# A novel k-mer set memory (KSM) motif representation improves regulatory variant prediction

**DOI:** 10.1101/130815

**Authors:** Yuchun Guo, Kevin Tian, Haoyang Zeng, Xiaoyun Guo, David Kenneth Gifford

## Abstract

The representation and discovery of transcription factor (TF) sequence binding specificities is critical for understanding gene regulatory networks and interpreting the impact of disease-associated non-coding genetic variants. We present a novel TF binding motif representation, the K-mer Set Memory (KSM), which consists of a set of aligned k-mers that are over-represented at TF binding sites, and a new method called KMAC for *de novo* discovery of KSMs. We find that KSMs more accurately predict in vivo binding sites than position weight matrix models (PWMs) and other more complex motif models across a large set of ChIP-seq experiments. KMAC also identifies correct motifs in more experiments than four state-of-the-art motif discovery methods. In addition, KSM derived features outperform both PWM and deep learning model derived sequence features in predicting differential regulatory activities of expression quantitative trait loci (eQTL) alleles. Finally, we have applied KMAC to 1488 ENCODE TF ChIP-seq datasets and created a public resource of KSM and PWM motifs. We expect that the KSM representation and KMAC method will be valuable in characterizing TF binding specificities and in interpreting the effects of non-coding genetic variations.

## INTRODUCTION

The binding of transcription factors (TFs) to specific short DNA sequences enables the precise control of gene expression in space and time. A TF binding motif is a short DNA sequence or sequences that a TF recognizes. We define the motif discovery task to be the identification of DNA sequences that are directly recognized by a TF, and thus are located at the site of binding where they mechanistically interact with a TF. Thus, our definition of a TF binding motif excludes co-factor motifs and other sequence features that are not immediately proximal to the site of TF binding.

Motifs are often used to identify preferential genome binding locations for a factor. Computational identification of TF binding sites are essential in deciphering transcriptional regulatory networks (Spellman et al. 1998; Lee et al. 2002; Kim and Park 2011). In addition, certain genetic variants associated with human diseases and phenotypic traits alter regulatory DNA sequences that are recognized by TFs (Maurano et al. 2012). Therefore, accurate TF binding motifs are critical to characterize TF binding differences between alleles and to identify the upstream regulators of non-coding variants (Claussnitzer et al. 2015). The advent of high throughput technologies such as ChIP-seq (Johnson et al. 2007) and protein binding microarrays (PBM)(Berger et al. 2006) have made a large amount of data available for the computation of in vivo and in vitro TF binding specificities. Computational methods for TF motif discovery remains an active and important area of investigation (Zambelli et al. 2012) and continues to inspire research into new approaches (Weirauch et al. 2013; Tompa et al. 2005).

Currently, there is no single standard for TF binding motif representation (Hughes 2011). The most widely used motif model is the position weight matrix (PWM) (Stormo 2000). However, the PWM model assumes that each base position contributes independently to binding probability and thus is unable to represent inter-base dependencies. Although PWM models provides a good approximation of protein-DNA interactions for many TFs (Benos et al. 2002; Zhao and Stormo 2011), dependencies between nucleotides at different positions in TF binding sites have been observed (Man and Stormo 2001; Bulyk et al. 2002; Berger et al. 2006; Maerkl and Quake 2007). In addition, a PWM is a highly compact and lossy representation. Therefore, in practice, PWMs fail to capture the full complexity of TF binding specificities in high throughput data.

K-mer based motif representations, which capture the exact bound sequences and thus preserve positional dependences if they exist, have been explored as alternatives to the PWM representation. Early work used individual over-represented k-mers to represent and discover TF binding motifs (van Helden et al. 1998; Tompa 1999). MotifCut connects k-mers into a graph and represents a motif as the maximum density subgraph, which is a set of k-mers that exhibit a large number of pairwise similarities (Fratkin et al. 2006). Recently, bags of k-mers (Ghandi et al. 2014) or clusters of k-mers (Setty and Leslie 2015) were used with binary classifiers for discriminating bound versus unbound sequences. However, the k-mer based representations of gkm-SVM (Ghandi et al. 2014) and seqGL (Setty and Leslie 2015) represent not only the DNA binding of a particular TF, but also other aspects such as chromatin accessibility and co-binding factor motifs. Thus gkm-SVM and seqGL fall outside of our definition of TF motif discovery. In addition, in these recent approaches, overlapping k-mers were implicitly assumed to be independent and were combined additively to score sequences. This independent assumption does not reflect the non-additive combinations of overlapping k-mers at a given site for binding, leading to an inaccurate representation of motifs.

More complex models accounting for positional dependencies have also been proposed, but they are rarely used in practice because they are computationally intensive and require more data to properly estimate the model’s parameters and may overfit if data are limited (MacIsaac and Fraenkel 2006; Zambelli et al. 2012). For example, The TF flexible model (TFFM) uses a hidden Markov model based framework to capture interdependencies of successive nucleotides and flexible length of the motif (Mathelier and Wasserman 2013). The sparse local inhomogeneous mixture (Slim) uses a soft feature selection approach to optimize the dependency structure and model parameters (Keilwagen and Grau 2015). Recently, deep neural network (deep learning) based approaches have been applied to predict TF binding with improved accuracy (Alipanahi et al. 2015; Zhou and Troyanskaya 2015). However, the distributed representation of deep learning models is more difficult to interpret mechanistically. We will compare deep learning based models with motif-based models for predicting the effect of non-coding genetic variants.

In addition, recent studies showed that proximal sequences flanking TF motifs may strongly affect the DNA shape and hence TF binding (Gordân et al. 2013; Levo and Segal 2014). Therefore, a motif model that preserves the base positional dependences in the motif and includes proximal flanking bases may improve the performance of PWM models and current k-mer based models.

In this paper, we present a novel motif representation that preserves the inter-position dependencies and includes the flanking k-mers, called K-mer Set Memory (KSM), and a de novo motif discovery method, K-Mer Alignment and Clustering (KMAC). A KSM consists of a set of aligned k-mers that are over-represented at factor binding sites and that can be combined nonadditively to accurately represent binding sites in new sequences. We show that KSM models predict in vivo TF binding more accurately than the PWM and the more sophisticated TFFM (Mathelier and Wasserman 2013) and Slim (Keilwagen and Grau 2015) models. In addition, predictive models based on KSM motifs outperforms those based on PWM motifs and deep learning derived sequence features in predicting differential regulatory activities of expression quantitative trait loci (eQTL) alleles. KMAC outperforms several state-of-the-art methods in correctly finding the known motifs from a large set of ChIP-seq data. Together, these results demonstrate that the KSM is a more accurate motif representation than the PWM and other representations for modeling TF binding and characterizing non-coding genetic variants.

## RESULTS

### The KSM motif representation

A TF’s *K-mer Set Motif* (KSM) is the set of overrepresented k-mers (gapped and ungapped words of length k) that are contained in the binding sites for the TF and have consistent offsets relative to the center of the binding sites (Figure 1A). The individual k-mers in a KSM are called *component k-mers*. A typical KSM may contain several hundred to several thousand component k-mers. Each component k-mer is annotated with a center offset and its presence/absence in each positive and negative training sequence. Unlike a PWM that assumes positional independence, KSM component k-mers are exactly matched to a query sequence being searched for a motif (Figure 1B). By requiring exact k-mer matches, a KSM preserves dependences among positions in the observed sequences. Long specific sequences are modeled when component k-mers overlap with each other (Figure 1A).

**Figure 1.**
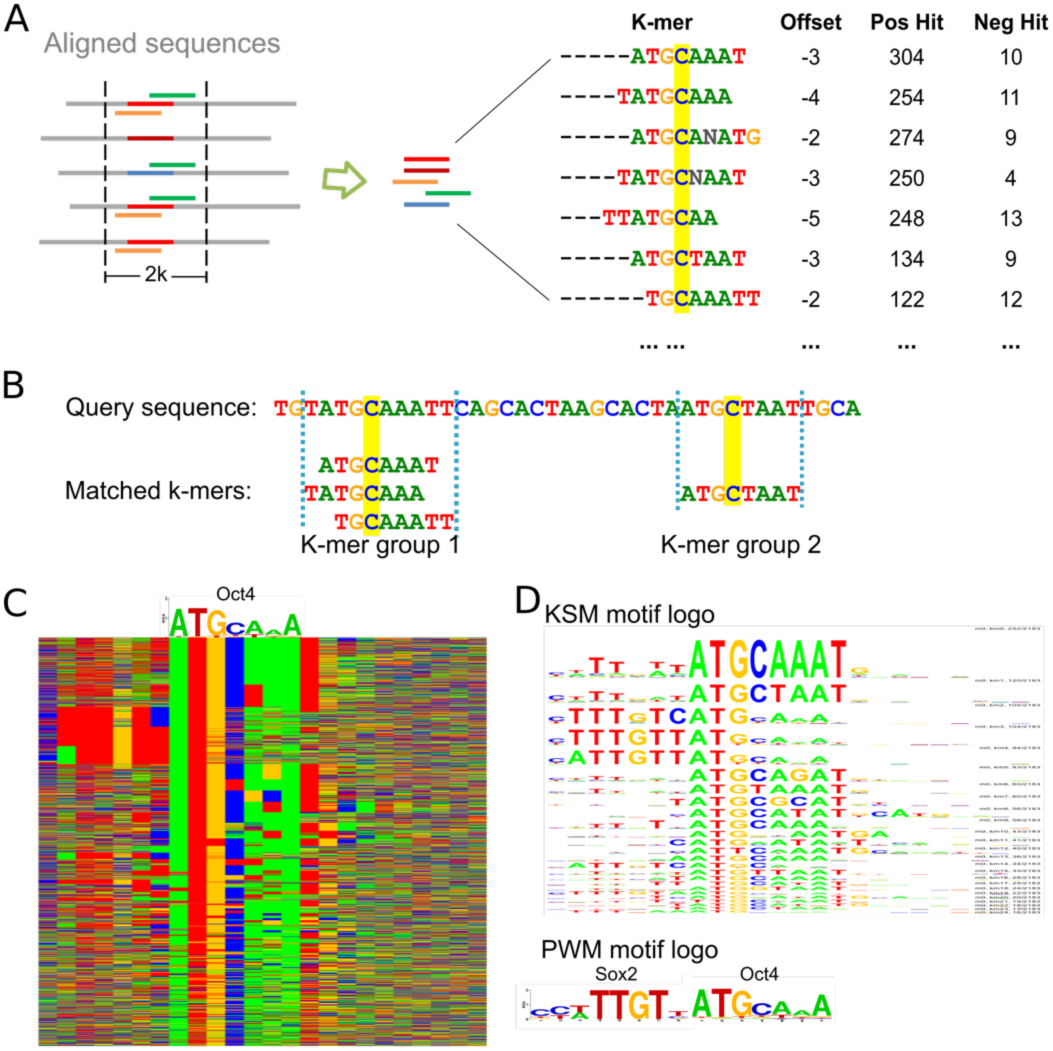
The KSM motif representation. (A) A KSM consists of a set of similar and consistently aligned component k-mers. The k-mers are extracted from a set of sequences aligned at the binding sites. Each k-mer has an offset that represent its relative position in the sequence alignment, and is associated with the IDs of the positive/ negative training sequences that contain the k-mer (total counts are shown). The base C highlighted in yellow represents the expected binding position. (B) An example of matching KSM motifs in a query sequence. (C) Color chart representation of 2183 sequences bound by Oct4 that match the Oct4 KSM motif. Each row represents a 23bp sequence. Rows are sorted by the KSM motif matches. Green, blue, yellow and red indicate A, C, G and T. An Oct4 PWM motif is shown above the sequences. (D) The KSM motif logo of Oct4 and the PWM logos of Sox2 and Oct4.

Each component k-mer is required to be overrepresented in the TF bound sequences (positive sequences) relative to the unbound sequences (negative sequences). We define the ***“sequence hit count”*** of a motif as the number of sequences containing the motif in the training sequence set, which is similar to the zero-or-one-per-sequence mode of MEME (Bailey and Elkan 1994). The over-representation of a component k-mer is evaluated by computing a hypergeometric p-value (HGP) (Barash et al. 2001):

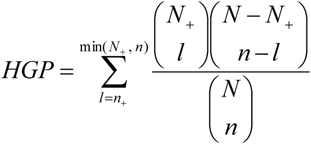

where *N* is the total number of positive and negative training sequences, *N*_+_ is the number of positive training sequences, *n* is the number of positive and negative training sequences containing the motif (positive and negative hit count), and *n*_+_ is the number of positive training sequences containing the motif (positive hit count). In this work, the component k-mers are required to have a HGP less than 10^−3^.

The offset of a component k-mer is defined as the offset of the first base of the k-mer relative to the expected binding center position, which is estimated during the motif discovery process (see below). For the Oct4 example, the expected binding position is the middle position of the binding site, i.e. base C (Figure 1A). When searching for a KSM motif in a query sequence (see next section), the offsets of the matched component k-mers can be used to align and group the k-mers that share the same expected binding positions into KSM motif instances called k-mer groups (Figure 1B).

A KSM’s representation of a large set of overlapping k-mers allows a KSM to capture the full complexity of TF binding specificities as well as the effect of the flanking bases, leading to a richer representation than the PWM and other consensus sequence representations (Stormo and Zhao 2010). For example, the Oct4 bound sequences also contains a Sox2 motif, which has been shown to have a strict spacing with Oct4 motif in mouse embryonic stem cells (Chew et al. 2005; Guo et al. 2012). The PWM motif learned from these sequences does not capture the existence of the Sox2 motif because the Sox2 motif only exists in a small subset of the sequences (Figure 1C). In contrast, the Oct4 KSM motif was able to capture the Sox2 motif through component k-mers such as TTTNTCATG and TTTGTCAT that overlap with both Oct4 and Sox2 motifs. To elucidate the complexities of the TF binding specificities, we compute a KSM motif logo to graphically represent the motif as a set of PWM motif logos that each summarize a component k-mer and its sequence context. From the KSM motif logo of Oct4, the existence of Sox2 motif can be easily observed (Figure 1D).

### KSM motif matching and scoring

To search for KSM motif instances in a query sequence, all of a KSM’s component k-mers are simultaneously searched using the Aho-Corasick algorithm for efficient multi-pattern search (Aho and Corasick 1975).

The k-mer matches in a query sequence are grouped into KSM motif instances based on their respective expected binding locations (Figure 1B), which are computed using the matched position of the k-mer and the KSM offset of the k-mer. We define a ***“k-mer group”*** (i.e. KSM motif instance) as the subset of component k-mers in the KSM model that occur in the query sequence and that are mapped to the same expected binding position on the sequence.

The hit count for a k-mer group cannot be obtained by simply summing the hit count of all the matching component k-mers, because the component k-mers are overlapping and a simple or weighted summation will not give an accurate count that re-capitulates the information in the training data. Thus the ***“k-mer group hit count”*** is defined as the number of all the training sequences that contain at least one of the matched k-mers in the k-mer group. In this formulation, overlapping k-mers are not combined additively as previous approaches (Ghandi et al. 2014; Setty and Leslie 2015), but in a non-additive manner that more accurately re-capitulates the contribution of these k-mers as a whole. Unlike the PWM motif instances, the KSM motif instances of the same motif may have different lengths because the length depends on the matched component k-mers and their relative positions.

The ***“KSM score”*** of a k-mer group is then defined as the odds ratio, which is a measure of association (Cornfield 1951; Edwards 1963):

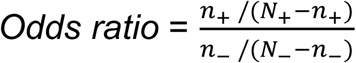

where *N*_+_ and *N*_-_ are the total numbers of positive and negative training sequences, respectively; *n*_+_ and *n*_-_ are the k-mer group positive and negative hit counts, respectively. To avoid divided-by-zero error, a small pseudo-count is added to the counts. If no component k-mer is matched in the query sequence, the KSM score of the sequence is 0.

### KMAC motif discovery

The K-Mer Alignment and Clustering (KMAC) method discovers both KSM and PWM motifs from a given set of positive (motif enriched) and negative sequences (motif depleted). If not provided, a negative sequence set is generated by randomly shuffling the positive sequences while preserving the di-nucleotide frequencies. KMAC can efficiently analyze the top 10,000 sequences from an assay and thus can learn weak signals. KMAC applies to sequences from in vivo TF ChIP-seq/ChIP-chip data or sequences of predicted elements from epigenomic data.

KMAC learns a KSM by aligning the positive sequences and computing the consistently aligned over-represented k-mers. KMAC uses values of k from 5 to 13 unless otherwise directed. For each value of k, KMAC discovers both KSM and PWM motifs as described below. All the motifs are then compared with each other and similar motifs are merged. Thus, the final list of motifs may be derived from different values of k, allowing KMAC to capture motifs with different lengths. KMAC motif discovery consists of four steps (Figure 2A):

**Figure 2.**
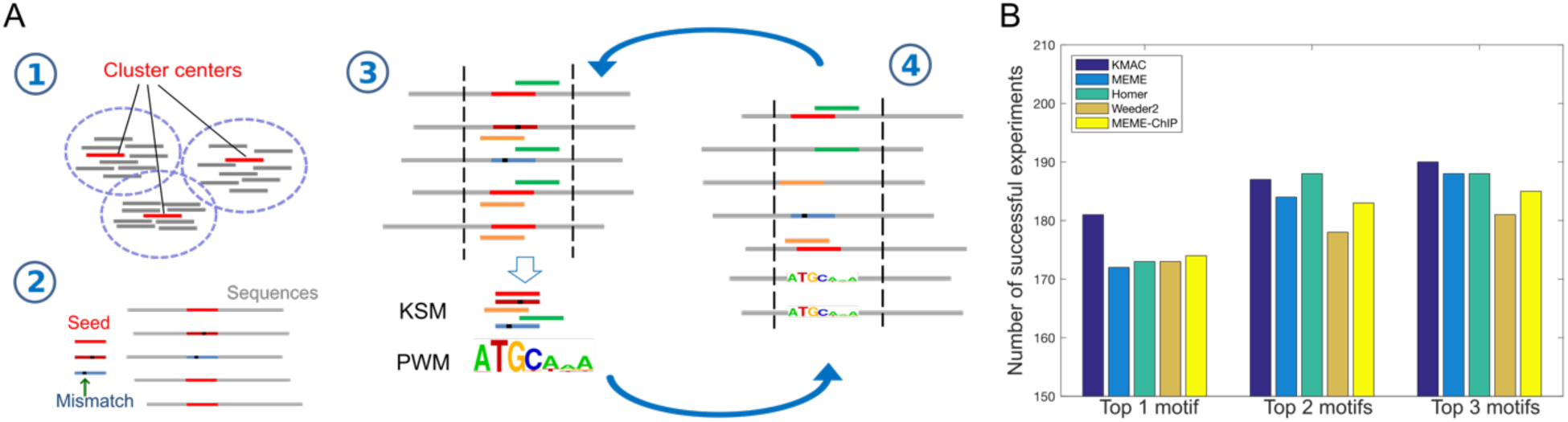
KMAC motif discovery outperforms other methods when detecting motifs in ChIP-seq data. (A) KMAC motif discovery schematic. Step 1: Over-represented k-mers with length k are clustered using density-based clustering. Bars represent the k-mers while red bars represent the cluster center exemplars. Step 2: A cluster center is used as a seed k-mer. The seed k-mer and k-mers with a one-base mismatch are used to match and align the sequences. Step 3: A pair of KSM and PWM motifs are extracted from the aligned sequences. Step 4: The KSM and PWM motifs are used to match and align the sequences. Step 3 and 4 are repeated until the significance of the motifs stops to improve. (B) The motif discovery performance of KMAC is compared to the motif discovery performance of various motif-finders on 209 ENCODE ChIP-seq experiments.

*Step 1:* KMAC selects a set of enriched k-mers and clusters them. K-mers with k exact bases and 0-4 contiguous gap bases are considered. The number of positive and negative sequences that contain instances of each possible k-mer are counted, treating each k-mer and its reverse complement as a single k-mer. A HGP is computed to evaluate the significance of enrichment for each k-mer. KMAC then clusters the enriched k-mers using a density-based clustering method (Rodriguez and Laio 2014). Levenshtein distance (Levenshtein 1966) is used to quantify the distance between two k-mers. KMAC then takes each of the top ranked k-mer cluster centers as the seed k-mers for the next phase. With the density-based clustering approach, a k-mer may belong to different k-mer clusters and thus contribute to different motifs, allowing KMAC to unbiasedly discover multiple motifs. This is in contrast to the typical mask-and-discover approach used by existing methods such as MEME (Bailey and Elkan 1994) and Homer (Heinz et al. 2010), which is biased in that the subsequently discovered motifs have a smaller sequence space.

*Step 2:* Each cluster center k-mer is used as a seed k-mer. This seed k-mer and similar k-mers with a one-base mismatch are used to initialize the KSM, which is then used to match and align the positive training sequences.

*Step 3*: A new KSM and its corresponding PWM motif are generated from a 2^*^k window around the middle of the seed k-mer using the alignment. To compute the offsets of the component k-mers, a reference position in the alignment, the expected binding position, is estimated as the median of the center positions of the aligned sequences.

*Step 4*: The KSM and PWM motifs are used to match and align the sequences. The KSM motif is first used to match the sequences and then the PWM motif is used to match the rest of the sequences. This allows KMAC to include more k-mers, especially at the initial iterations when the KSM consists only a few component k-mers. If multiple motif matches are found in a sequence, the match with the highest score is used.

Step 3 and 4 are repeated alternately until the significance of the motifs stops to improve. The significance of a motif is evaluated as the sum of partial area under receiver operating characteristic (pAUROC) (up to a false positive rate of 0.1, fpr<=0.1) scores of the KSM and PWM motifs (Ma et al. 2013; McClish 1989). We choose the pAUROC because typically only the area at false positive rate less than or equal to 0.1 is of interest for realistic motif matching.

Finally, all the discovered motifs are ranked by the sum of KSM and PWM pAUROC scores.

### KMAC outperforms other motif discovery methods in discovering known DNA-binding motifs

We tested KMAC’s ability to discover biologically relevant DNA binding motifs in data from the ENCODE project (The ENCODE Project Consortium 2012). We used a set of 209 TF ChIP-seq experiments and associated controls comprising 78 distinct TFs that were profiled in one or more cell lines by the ENCODE project and for which validated DNA binding motifs exist in public databases (Weirauch et al. 2014). We chose this large collection of experiments because we expected that they would be representative of the typical range of ChIP-seq data noise and sequencing depth. We used KMAC and four state-of-the-art methods, MEME (Bailey and Elkan 1994), MEME-chip(Machanick and Bailey 2011), Homer (Heinz et al. 2010), and Weeder2 (Zambelli et al. 2014) to analyze DNA sequences derived from these ChIP-seq data. The most significant motifs from each analysis were compared to corresponding known binding motifs of the same TFs using STAMP (Mahony et al. 2007). We found that KMAC outperforms other methods in rediscovering the known motifs in the public database cisBP (Weirauch et al. 2014) (Figure 2B). When allowing each method to make multiple motif predictions, KMAC performs better than or equal to the other methods. In addition, the running time of KMAC is similar to that of Weeder2, and is much faster (about 4x-30x) than the other methods (Table S1).

### KSMs outperform PWMs in predicting in vivo TF binding

We compared the performance of KSMs versus PWMs in predicting in vivo TF binding using the ENCODE ChIP-seq datasets. We found that the KSM outperforms the PWM in discriminating TF bound sequences (positive sequences) from randomly generated sequences and unbound genomic sequences near the binding sites (negative sequences).

First, we trained KSM and PWM motifs using a subset of TF GABP-bound sequences in human K562 cells, and used the motif scores to discriminate held-out GABP-bound sequences from negative randomly shuffled sequences. We found that the KSM outperforms three PWMs learned by KMAC, MEME, and Homer, respectively, from the same set of sequences (Figure 3A). To understand why the KSM performs better than the PWM, we next studied the sequences and scores of the GABP motif matches. We found that for the same PWM motif match scores, the KSM scores of the matches in the positive sequences are generally higher than the KSM scores of those in the negative sequences (Figure 3B). The higher KSM scores in the positive sequences are contributed by the overlapping k-mers that are often present in the positive sequences but are less present in the negative sequences. These results are consistent with the observation that the length of the motif matches in the positive sequences are longer than the length in the negative sequences (Figure S1). Therefore, the KSM is able to use the flanking sequences to further discriminate real bound sequences from the random sequences when they have identical PWM matches. In addition, we found cases that some sites in the negative sequences are scored highly by the PWM but not by the KSM. For example, CACTT<u>G</u>CGG is only one base different from the consensus sequence CACTTCCGG and has a PWM score of 6.67, which is about 60% of the maximum PWM score for GABP motif. However, CACTTGCGG does not occur in the entire GABP bound sequence set, suggesting this single base difference cannot be tolerated by GABP. The KSM score for the CACTTGCGG site is 0 because it has no exact match to the KSM. This highlights the limitation of the positional independence assumption of the PWM representation and that the KSM is able to overcome this limitation.

**Figure 3.**
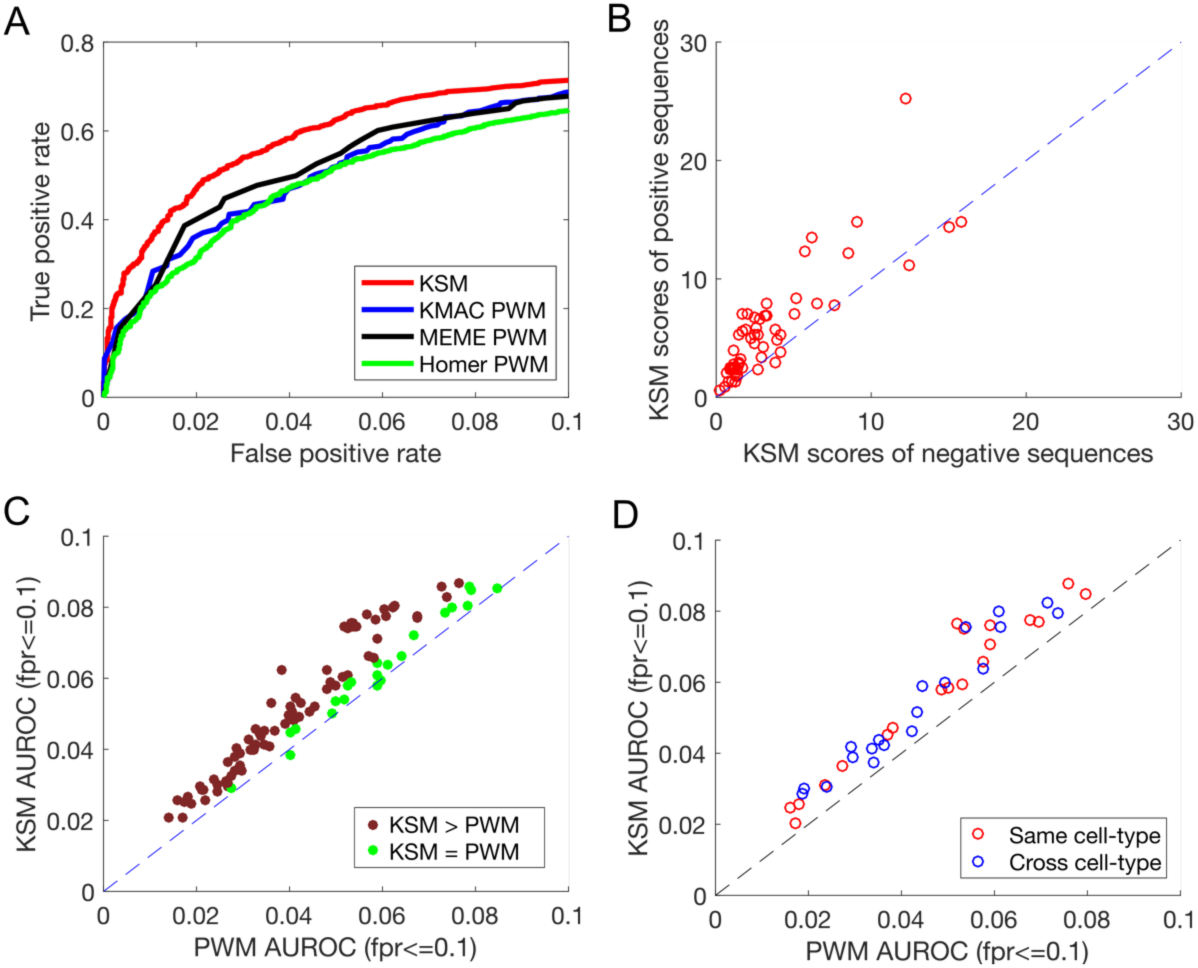
KSM outperforms PWM in predicting in vivo TF binding in held-out data. (A) The partial ROC performance of KSM, KMAC PWM, MEME PWM, and Homer PWM for predicting ChIP-seq binding of GABP in K562 cells. (B) Scatter plot comparing the mean KSM scores of positive sequences and mean KSM scores of negative sequences that corresponds to the same PWM scores in the K562 GABP dataset. Each point represents a set of sequences that have the same PWM score. (C) Scatter plot comparing the median partial AUROC (fpr<=0.1) values of KSM and PWM for predicting ChIP-seq binding for 43 TFs. (D) Similar to (C), but comparing KSM and PWM in the same cell type (red) or across cell type (blue) in 19 TFs.

We then extended the comparison between the KSM and the PWM to 103 datasets where the correct primary motifs were found and that have sufficient number of binding sites. Here the KSM and PWM motifs were both learned from the same KMAC motif discovery runs to ensure that the performance differences are from the motif representation but not from the motif discovery procedures. Similar to previous work (Mathelier and Wasserman 2013), we compare the performance between two methods by computing the score ratios between the methods on the same datasets. Two methods are considered performing differently if the score ratio is less than 0.95. In 94 out of 103 experiments, the KSMs perform better than the PWMs in predicting TF binding in held-out data, while the PWMs do not perform better in any of the experiments (Figure 3C). Across all the datasets, the KSM representation significantly outperforms the PWM representation (p=1.53e-18, paired Wilcoxon signed rank test). We also tested using flanking sequences as negative sequences, and obtained similar results (p=4.44e-15, paired Wilcoxon signed rank test) (Figure S2).

We also found that a KSM does not overfit the training data and is able to generalize across cell types. Because a KSM consists of hundreds to a few thousand k-mers, one legitimate concern is that it may overfit the training data. Overfitting would cause good performance on the training cell type but poor performance on a new cell type. To address this concern, we conduct a crosscell-type analysis. For 19 unique TFs that are both profiled in different cell types by the ENCODE project, including a diverse list of CTCF, NRSF, YY1, USF, PU.1, E2F6, c-Jun, ETS1, etc., we trained KSM and PWM motifs from one cell type (K562) and predicted binding for another cell type (GM12878 or H1-hESC). We found that KSMs significantly outperformed PWMs in the cross-cell-type prediction (p=0.000132 for same cell type, and p=0.000132 for cross cell-type preditions, paired Wilcoxon signed rank test) (Figure 3D). The KSM predictions across the cell types perform similarly to the KSM predictions in the same cell type (p>0.05, paired Wilcoxon signed rank test).

Taken together, these results suggest that the KSM is a more accurate motif representation than the PWM model.

### KSM outperforms complex motif models in predicting in vivo TF binding

We next compared the KSM representation with two complex motif models that have been shown to be more accurate than the PWM model. The TF flexible model (TFFM) is a hidden Markov model based framework that captures interdependencies of successive nucleotides and flexible length of the motif (Mathelier and Wasserman 2013). The sparse local inhomogeneous mixture (Slim) uses a soft feature selection approach to optimize the dependency structure and model parameters (Keilwagen and Grau 2015). We trained TFFM and Slim models on the same subset of sequences as the KSMs and used the motif scores to predict on the rest of the sequences. The KSMs perform better than the TFFMs in predicting TF binding in 39 experiments, worse in 4 experiments, and similarly in 60 experiments (Figure 4A). Across all the datasets, the KSM significantly outperforms the TFFM representation (p=2.03e-5, paired Wilcoxon signed rank test). Similarly, the KSMs perform better than slim in predicting TF binding in 43 experiments, worse in 22 experiments, and similarly in 38 experiments (Figure 4A). Across all the datasets, the KSM significantly outperforms the Slim representation (p=1.79e-3, paired Wilcoxon signed rank test). In addition, the motif scanning time of KMAC is only 2-3 times of the PWM scanning time, and is much less (about 20x-80x) than that of the Slim and TFFM models (Table S2).

**Figure 4.**
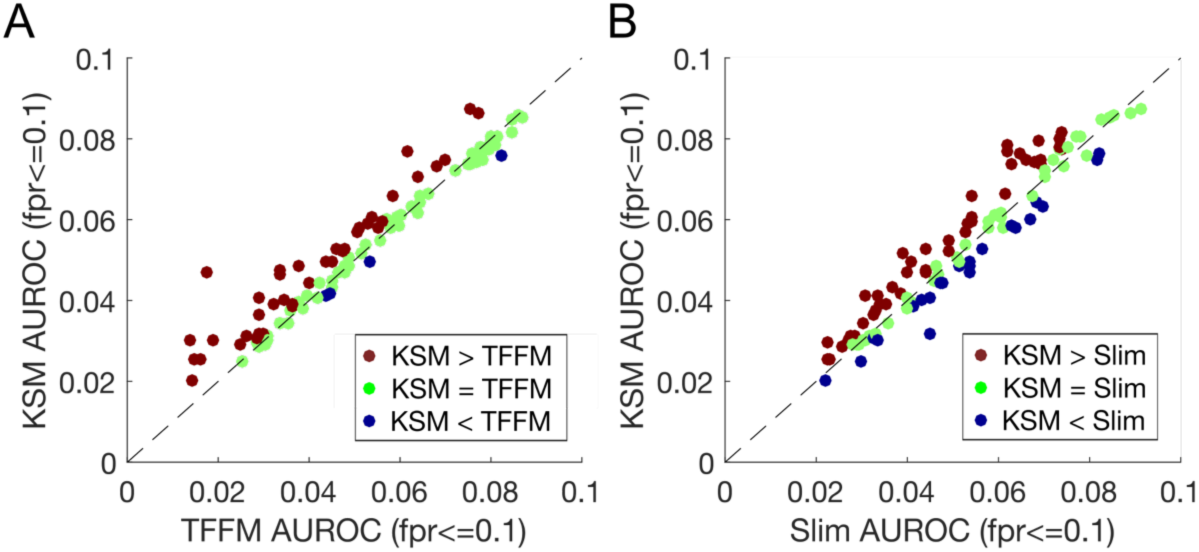
KSM outperforms complex motif models in predicting in vivo TF binding. (A) Scatter plot comparing the mean partial AUROC (fpr<=0.1) values of KSM and TFFM for predicting in vivo binding in 103 TF ChIP-seq experiments. Each point represents a ChIP-seq dataset. (B) Similar to (A), but comparing KSM and Slim.

In summary, the KSM is more accurate at discriminating TF bound sequences from randomly generated sequences than the conventional PWM and the more sophisticated TFFM and Slim motif representations, suggesting that the KSM is a more precise motif representation.

### Incorporating DNA shape features improves in vivo TF binding prediction

We next investigate whether incorporating DNA shape features with KSM can further improve the in vivo TF binding prediction. DNA shape features inside the motif and at the flanking bases of the motifs have been shown to improve TF binding prediction with the PWM and the TFFM (Mathelier et al. 2016). Because the KSM may capture DNA shape information implicitly through the k-mers that it contains, it is of interest to know how much the explicit DNA shape information can further improve the prediction accuracy of the KSM motifs and how does the improvement compare with the improvement from incorporating DNA shape information with the PWM.

Similar to previous work (Mathelier et al. 2016), we trained gradient boosting classifiers (Friedman 2001) to predict TF binding with four different set of features: KSM score, KSM score + DNA shape, PWM score, and PWM score + DNA shape. We constructed datasets that consist of 101bp positive sequences around the 10,000 top-ranked ChIP-seq binding events and 101bp negative randomly shuffled sequences. We accessed the performance of the classifiers through the average area under precision-recall curve (AUPRC) metrics from 10-fold cross validation (CV). For each CV set, we learned KSM and PWM motifs from the training set, verified that the motifs matched known motifs in the public cisBP database (Weirauch et al. 2014), and then selected only the sequences that contains both KSM and PWM motif matches to construct balanced training and testing sets for the classifiers. For this analysis, all the positive and negative sequences contain motifs and thus the prediction task is more challenging than using un-selected sequences. The KSM and PWM motif scores and DNA shape features were then used to construct four types of features.

We found that incorporating DNA shape features (110 features) with a KSM or PWM motif score (1 feature) improves the in vivo TF binding prediction across all the datasets we tested (Figure 5A, S3, and S4), consistent with previous findings (Mathelier et al. 2016). In addition, incorporating shuffled DNA shape features does not improve the prediction upon the KSM/PWM motif score alone (Figure S5), suggesting that information encoded in the shape features contributes to the prediction. Remarkably, factors such as CTCF, MafK and MafF achieve high accuracy (AUPRC=0.965~0.975) with the KSM/PWM and DNA shape (31bp) combined features.

**Figure 5.**
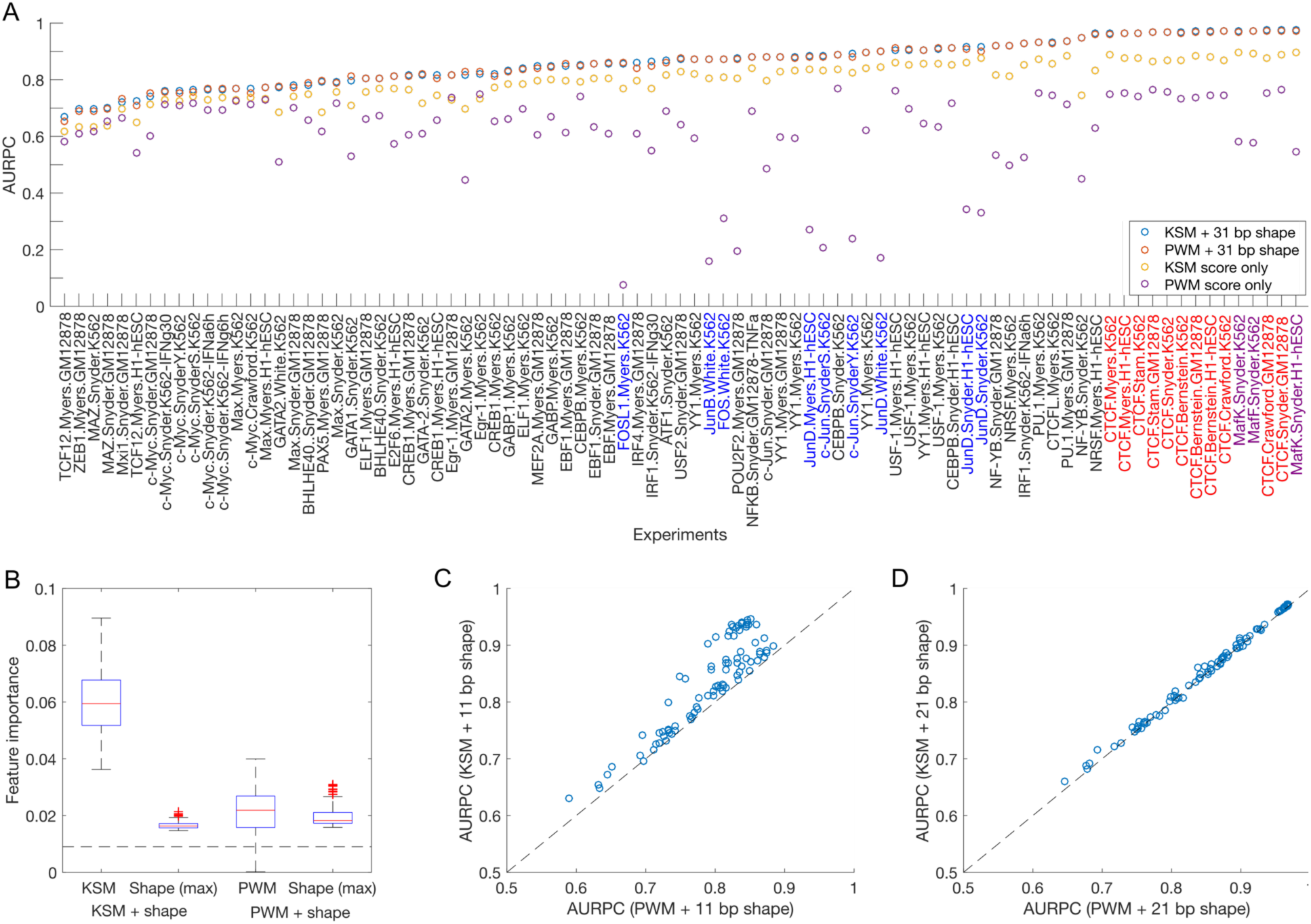
Incorporating DNA shape features improves in vivo TF binding prediction. (A) AUPRC performance comparison among models trained with various set of features: KSM, PWM, KSM + DNA shape, and PWM + DNA shape. (B) Feature importance measures from the gradient boosting models across all the ChIP-seq datasets show that the KSMs contribute more to the prediction than the PWMs. Horizontal dashed line indicates the average feature importance for 1+110 features in the motif + 31bp DNA shape models. (C) Scatter plot comparing the AUPRC performance between KSM + 11bp DNA shape and PWM + 11bp DNA shape features, each point is a ChIP-seq dataset. (D) Similar to (C), but with 21bp DNA shape features.

Without the DNA shape features, KSM score alone performs much better than PWM score alone in most of the experiments (Figure 5A), especially for AP-1 factors such as c-Jun, c-Fos, JunB, JunD, and FOSL1. Incorporating the DNA shape features appears to make up most of the differences between the KSM and PWM motifs. Although KSM + shape (31bp) features results in classifiers with slightly higher AUPRC scores than PWM + shape (31bp) features overall (p=0.005, paired Wilcoxon signed rank test), the absolute differences are very small (Figure 5A). With the DNA shape features, the performance gains of the KSMs are much smaller than those of the PWMs. This finding is consistent with further analysis on the feature importance metrics from the classifier with motif + shape features. In the KSM + shape models for all the datasets, the feature importance of the KSM score is much higher than the maximum importance value of all the shape features (p=5.46e-16, paired Wilcoxon signed rank test) (Figure 5B), suggesting that the KSM consistently contributes more than any single shape feature to predict TF binding. In contrast, in the PWM + shape models, the feature importance of the PWM score is comparable to (p>0.05, paired Wilcoxon signed rank test) and in some cases lower than the maximum importance value of all the shape features (Figure 5B). These results suggest that the PWM suffers from information loss and relies more on the shape features to predict TF binding, and the KSMs have captured additional sequence information, including some information represented by the DNA shape features.

To further understand the effect of DNA shape features from the bases inside the motif and those from the flanking bases, we also tested DNA shape features from 21bp and 11bp sequences around the motif positions. For most of the TFs we tested, the 11bp shape features are sufficient to cover the motif cores, and the 21bp shape features cover some additional flanking bases. With the 21bp shape features, KSM + shape models perform similarly to the PWM + shape models (Figure 5D). In contrast, with the 11bp shape features, KSM + shape models significantly outperform the PWM + shape models (p=5.63e-16, paired Wilcoxon signed rank test) (Figure 5C). These results suggest that the shape information from the motif cores is not sufficient to make up the differences between the KSM and the PWM. The additional information captured by the KSM may partly come from the k-mers that covers the flanking bases. For most of the datasets, for both the KSM and the PWM, incorporating DNA shape information from a larger region around the motif consistently results in better prediction (Figure S3 and S4). However, 31 bp shape features only performs slightly better than the 21bp shape features (Figure S3 and S4), suggesting shape features further than 21bp may not add much additional benefits.

### KSM enables accurate prediction of causal regulatory variants

With the superior performance of the KSM representation on predicting in vivo TF binding, we next tested whether sequence features derived from KSM motifs would enable more accurate prediction of the effects of non-coding genetic variants on the activities of the regulatory sequences that harbor the genetic variants.

We used an ensemble model (Zeng et al. 2017) that included KSM motif features from 87 TF ChIP-seq datasets and deep learning based features to achieve the best performance in “eQTL-causal SNPs” open challenge (Kreimer et al. 2017) in the Fourth Critical Assessment of Genome Interpretation (CAGI 4). The challenge was to predict the experimental results of thousands of regulatory elements that contains eQTL alleles (reference and alternative) from a massively parallel reporter assay in GM12878 cells (Tewhey et al. 2016).

Here, we used the same computational framework, a LASSO regression model to predict reporter expression of the reference and alternative alleles and an ensemble model to classify whether the two alleles have different regulatory activities (Zeng et al. 2017), to evaluate the performance of different types of sequence features. We constructed KSM motif features and PWM features from motifs discovered by MEME and Homer, respectively, from 209 TF ChIP-seq datasets (The ENCODE Project Consortium 2012). The performance of the predictions using different set of sequence features was evaluated using AUPRC and AUROC. We found that the KSM features (AUPRC=0.479, AUROC=0.668) outperform Homer PWM (AUPRC=0.434, AUROC=0.629) and MEME PWM (AUPRC=0.408, AUROC=0.619) in predicting differential reporter expression between the two alleles (Figure 6A and S6A). We next compared KSM motif features with the features derived from DeepBind (Alipanahi et al. 2015), a deep learning model trained on 927 TF ChIP-seq datasets, and DeepSEA (Zhou and Troyanskaya 2015), a deep learning model trained on 919 epigenomic datasets. We found that KSM features outperform DeepBind (AUPRC=0.432, AUROC=0.608) and DeepSEA features (AUPRC=0.396, AUROC=0.628) in predicting differential reporter expression between the two alleles (Figure 6B and S6B). In addition, the KSM features offer better interpretability than deep learning features because the predictive KSM features are directly linked to their corresponding TFs. The combined KSM and DeepBind features achieved the best AUPRC (0.483), outperforming the KSM or DeepBind features alone, although the AUROC (0.647) of the combined features is worse than that of the KSM. The combined KSM and DeepBind features or KSM features alone both outperform all the CAGI 4 methods that use features such as PWMs, k-mers, epigenomic signals, chromatin state annotations, and evolutionary conservation (Kreimer et al. 2017). These results highlight the value of accurate motif models in the characterization of non-coding variants.

**Figure 6.**
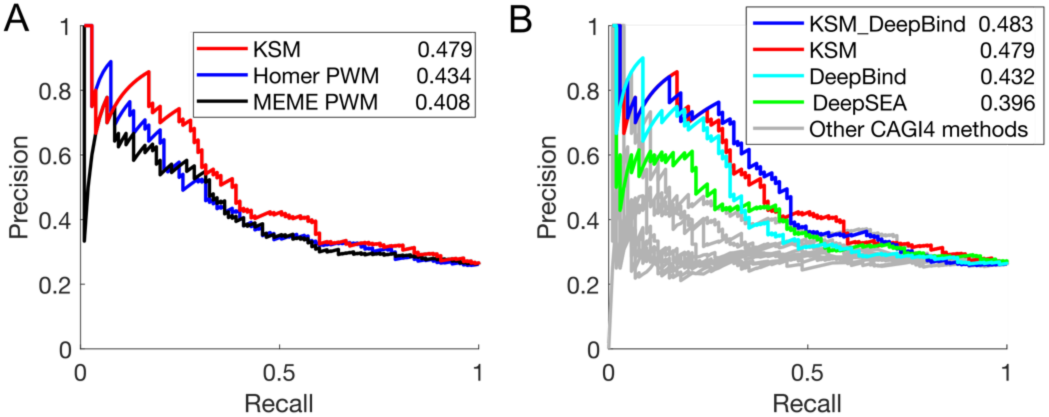
KSMs predicts allele-specific differences in regulatory activity better than PWMs and deep learning derived features. (A) PRC performance of KSM and PWM motif representations in predicting differential regulatory activities of eQTL alleles. The numeric values in the legend are the AUPRC values. (B) Similar to (A), KSM, DeepBind, DeepSEA derived features and other CAGI 4 open challenge methods.

### A new public resource of KSM and PWM motifs

Finally, we have created a public resource of KSM and PWM motifs by applying KMAC to 1488 ENCODE transcription factor ChIP-seq datasets.

## DISCUSSION

We have demonstrated that k-mer set memory (KSM) representations are better at predicting transcription factor in vivo binding than PWMs and the more complex TFFM and Slim models. In addition, sequence features derived from KSMs outperform those derived from PWMs and deep learning models for predicting the effect of non-coding genetic variants. Given that most computational methods that involves TF binding motifs use the PWM representation, the accuracy gain from replacing the PWM with the KSM will likely be wide spread.

A KSM represents factor binding specificity as a set of aligned k-mers that are found to be over-represented at factor binding sites. An important feature of the KSM is that it captures the relative positions among the k-mers, thus allowing overlapping k-mers to be assembled into k-mer groups for accurate identification and scoring of motif instances. We showed that, with contribution from the overlapping k-mers, the KSM gives TF bound sequences higher scores than the random sequences when they have the same PWM score, highlighting the value of positional information among the k-mers for recapitulating in vivo TF binding.

The KSM is a representation for the DNA binding motif of a single TF. To increase the probability that a KSM represents the sequence mechanistically associated with a particular TF, KMAC uses a narrow window around binding sites to extract component k-mers and requires the component k-mers to be aligned with each other. Thus, the KSM is different from and not directly comparable to methods for alternate tasks that use k-mers associated with multiple TF motifs in machine learning models (Ghandi et al. 2014; Setty and Leslie 2015). It will be interesting to build learning models with multiple KSM motifs learned from ChIP-seq or DNase-seq data and compare with the published k-mer-based learning methods.

DNA shape at the flanking bases and the motif cores have been shown to important for TF binding (Abe et al. 2015; Levo and Segal 2014; Slattery et al. 2011). We showed that incorporating DNA shape information further improves upon KSM motifs for predicting in vivo TF binding. The DNA shape features used in our analysis were derived from 5-mer sequences (Chiu et al. 2016). Therefore, the improvement from incorporating DNA shape features may come from the information in the 5-mer sequences that are not necessary shape-related information (Zhou et al. 2015). The 5-mer or shorter sequences may not be fully captured by the current KSM models that typically use longer k-mers (k=7~12). In addition, our analysis suggested that the flanking k-mers in the KSM representation contribute to the advantage of the KSM over the PWM; and incorporating DNA shape features from the motif-flanking bases further out (e.g. 21bp) showed better prediction accuracy. These results suggest that the KSM representation may be further improved by integrating the flanking bases, DNA shape information, or other short k-mer representations. Although DNA shape information may augment the prediction accuracy of a motif, in practice, it is challenging to fully integrate the DNA shape information because of its size. For example, for a single motif feature, an additional 110 features are needed to encode the 31bp first-order DNA shape information, and the feature size doubles if second-order DNA shapes are considered. For a comprehensive model that includes hundreds or more motifs, the large feature set size may lead to a high computation load and may cause the model to overfit. Therefore, a more comprehensive motif model that integrates the DNA shape information around the motif could be valuable.

Genome-wide association studies (GWAS) have made tremendous progress in linking numerous single nucleotide polymorphisms (SNPs) to human traits and diseases. However, finding the causal genetic variants has been challenging because the lead GWAS SNPs are in linkage disequilibrium with nearby SNPs and the majority of GWAS loci are in non-coding regions (Maurano et al. 2012; Schaub et al. 2012). Computational approaches that identify TF binding altering genetic variants are important for meeting this challenge (Mathelier et al. 2015). The KSM motif representation and the KMAC motif discovery method enables more accurate characterization and discovery of TF binding motifs. Our results show that the KSM motif features outperform features derived from deep learning model in predicting the effect of non-coding genetic variants, suggesting that accurate and interpretable motif features may be more appropriate for characterizing non-coding genetic variants than the deep learning features. With large scale efforts such as the ENCODE project (The ENCODE Project Consortium 2012) profiling hundreds of TFs in diverse cellular conditions, a more comprehensive catalog of TF binding sites is now available for training new computational models. We expect that the KSM representation and KMAC method will be valuable in characterizing TF binding specificities and in interpreting the effects of non-coding genetic variations.

## METHODS

### ChIP-seq datasets and TF binding motifs

209 TF ChIP-seq datasets (from three ENCODE tier 1 cell types, K562, GM12878, and H1-hESC cells) that have known motifs in public databases were downloaded from the ENCODE project website (The ENCODE Project Consortium 2012). TF binding motifs (PWMs) were downloaded from cisBP database (Homo_sapiens_2015_02_05)(Weirauch et al. 2014), which includes motif from the TRANSFAC (Matys et al. 2003), JASPAR (Sandelin et al. 2004), and Uniprobe (Berger et al. 2006) databases.

### Motif discovery performance comparison

For the 209 ENCODE ChIP-seq data, KMAC and four other state-of-the-art *de novo* motif discovery methods, MEME v4.11 (Bailey and Elkan 1994), MEME-chip v4.11 (Machanick and Bailey 2011), Weeder 2.0 (Zambelli et al. 2014), and Homer (Heinz et al. 2010), were applied to discover motifs independently. From the top 1000 peaks of each dataset, 100bp sequences centered on the peak summits were extracted, as suggested by the MEME Suite’s documentation based on the typical resolution of ChIP-seq peaks. MEME was run with options “-dna -nmotifs 9-revcomp”, MEME-chip was run with options “-dna -norand -meme-nmotifs 5 -meme-maxsize 1000000 -dreme-m 5 -spamo-skip -fimo-skip”, and Weeder2 was run with options “-O HS - chipseq”. All other parameters were the defaults specified by the authors.

Discovered motifs (PWMs) were compared to known motifs in the public database cisBP (Weirauch et al. 2014) using STAMP (Mahony et al. 2007). For KMAC, the PWM motifs discovered were used for comparison. A motif with E-value less than 1e-5 was considered a match. For each program, we counted the number of datasets that had a motif matching at least one known motif of that TF. In some cases, the correct motifs were not matched by the first motif that a method outputs, but by the second or later motifs. Therefore, we compared the motif-finding performance using the top 1, top 2, and top 3 motifs.

### TF in vivo binding prediction performance comparison

We compared KSM, PWM, Slim (Keilwagen and Grau 2015) and TFFM (Mathelier and Wasserman 2013) motif models in predicting in vivo TF ChIP-seq binding sites. For each set of bound sequences from a TF ChIP-seq experiment (positive sequences), we generated random shuffled sequences by preserving di-nucleotide frequencies (shuffled negative sequences). We also generate an alternative set of negative sequences by taking the genomic sequences 200bp away from the TF binding site (flanking negative sequences).

We first discover motifs from randomly subsampled 5000 positive sequences (training set) using KMAC, Homer, MEME, the Jstacs library for Slim (Keilwagen and Grau 2015), and the Python TFFM framework (Mathelier and Wasserman 2013). For Slim, additional shuffled negative sequences with signal=0 were provided for motif discovery. For TFFM, two kinds of models (FIRST_ORDER and DETAILED) were constructed by using the primary PWM motifs discovered by MEME for initialization. The results from two TFFM models were similar. We report only results from the DETAILED model.

For each method, the motif scores of the top ranking primary motif are then used to discriminate 5000 held-out positive and negative sequences (test set). In order to compare performance across multiple motif representations, we used 113 datasets where the primary motifs discovered by all the methods/representations match a known motif for the same TF in the public database cisBP (Weirauch et al. 2014). For the different motif representations discovered from the same set of sequences, their performance in predicting ChIP-seq TF binding sites on the held-out data was evaluated using a partial AUROC (McClish 1989) up to a false positive rate of 0.1, which typically falls in the range of realistic motif matches. We repeated this procedure five times and used the mean partial AUROC score of each ChIP-seq experiment or each TF to compare performance.

We assessed the significance for the improvement of predictive power when comparing two models using the Wilcoxon signed rank tests. The function signrank() In MATLAB software (MATLAB and Statistics Toolbox Release 2016b, The MathWorks, Inc., Natick, Massachusetts, United States) was used.

### TF binding prediction with motif and DNA shape

From the ENCODE ChIP-seq datasets that contains at least 10,000 binding events, we construct datasets that include 101bp positive sequences around the 10,000 top-ranked binding sites of each ChIP-seq dataset and 101bp negative sequences generated by randomly shuffling the positive sequences while preserving the di-nucleotide frequencies. We accessed the performance of the classifiers through 10-fold cross validation (CV).

For each CV dataset (9,000 training sequences and 1,000 testing sequences), we learned KSM, PWM motifs from training set using KMAC and use the top-ranked motif for the subsequent analysis. Discovered PWM motifs were compared to known motifs in the public database cisBP (Weirauch et al. 2014) using STAMP (Mahony et al. 2007). To avoid the differences due to motif discovery, we used 87 experiments that all 10 CV sets produced the correct primary motif of the corresponding TF. Then we constructed motif score and DNA shape features by scanning for both the KSM and PWM motifs on the sequences, and generated DNA shape features from sequences around the motif matches. To ensure that the sequences contain motifs and sufficient sequence to generate DNA shape features, we used a selection criteria that require the sequences (both positive and negative, training and testing) to contain both KSM and PWM motif matches and have at least 31bp sequence around the motif match positions. Because negative sequences typically contain less motif matches than positive sequences, we generated multiple set of negative sequences with different random seeds such that the positive and negative sequence set that meet the selection criteria have the same number of sequences (i.e. balanced dataset).

DNA shape features were generated using the DNAshapeR R/Bioconductor package (Chiu et al. 2016). For this study, we only used the first-order shape features: HelT, MGW, ProT, and Roll. We generated 3 different set of DNA shape features from 31bp, 21bp, and 11bp sequences around the motif (KSM or PWM) positions.

We used the GradientBoostingClassifier in the Python scikit-learn module (Pedregosa et al. 2011) (version 0.18) to train, and apply gradient boosting classifiers to DNA sequences. Features used in the classifiers were vectors composed of motif score (KSM or PWM) (size=1), and the four first-order DNA shape features at each nucleotide (size=~4n). The feature matrix was standardized column-wise. The parameters for the classifier were: learning_rate=0.1, max_depth=4, n_estimators=500. We tested other parameter settings and the performances are similar. The feature importance value for each feature were obtained from the GradientBoostingClassifier method. The feature values were normalized to sum to 1.

The probabilities generated by gradient boosting classifiers were used to compute AUPRC performance metrics using the MATLAB software (MATLAB and Statistics Toolbox Release 2016b, The MathWorks, Inc., Natick, Massachusetts, United States). The mean value of the AURRC from the 10 CV sets of each dataset is presented.

### Predicting the effect of regulatory variants

We used the EnsembleExpr <u>(https://github.com/gifford-lab/EnsembleExpr/)</u> computational framework as described in (Zeng et al., 2016). Briefly, sequence features were generated by taking the maximum motif score of each motif on the training and testing sequences. LASSO regression models were trained to predict the reporter expression levels for each allele, and an ensemble of binary classification models with regularization tuned by cross-validation was trained to predict whether the two alleles have different expression levels. In this work, 5 set of sequence features were derived from KSM motifs, MEME PWM motif, and Homer PWM motifs learned from 209 ENCODE TF ChIP-seq datasets, and from the pre-trained DeepBind (Alipanahi et al. 2015) and DeepSEA model (Zhou and Troyanskaya 2015).

## Software availability

The KSM and KMAC free software can be downloaded from <u>(http://groups.csail.mit.edu/cgs/gem/kmac/).</u>

## AUTHOR CONTRIBUTIONS

YG conceived the project. YG and DKG designed the analysis. YG developed the KSM and KMAC methods. YG coordinated the analysis. YG, KT, HZ, and XG performed the analysis and interpreted results. YG and DKG wrote the manuscript.

## ACKNOWLEDGMENTS

We thank Jens Keilwagen for providing suggestions and codes for training and using Slim model. This work was supported by National Institutes of Health (grant 1U01HG007037-01 to D.K.G).

## SUPPLEMENTAL INFORMATION

Supplemental Information includes two tables and six figures can be found with this article online

